# Genomic and phenotypic comparison of environmental and patient-derived isolates of *Pseudomonas aeruginosa* suggest that antimicrobial resistance is rare within the environment

**DOI:** 10.1101/663674

**Authors:** Kay A Ramsay, Samuel J T Wardell, Wayne M Patrick, Ben Brockway, David W Reid, Craig Winstanley, Scott C Bell, Iain L Lamont

## Abstract

Patient-derived isolates of the opportunistic pathogen *Pseudomonas aeruginosa* are frequently resistant to antibiotics due to the presence of sequence variants in resistance-associated genes. However, the frequency of antibiotic resistance and of resistance-associated sequence variants in environmental isolates of *P. aeruginosa* has not been well studied. Antimicrobial susceptibility testing (ciprofloxacin, ceftazidime, meropenem, tobramycin) of environmental (n=50) and cystic fibrosis (n=42) *P. aeruginosa* isolates was carried out. Following whole genome sequencing of all isolates, twenty-five resistance-associated genes were analysed for the presence of likely function-altering sequence variants. Environmental isolates were susceptible to all antibiotics with one exception, whereas patient-derived isolates had significant frequencies of resistance to each antibiotic and a greater number of likely resistance-associated genetic variants. These findings indicate that the natural environment does not act as a reservoir of antibiotic-resistant *P. aeruginosa*, supporting a model in which antibiotic susceptible environmental bacteria infect patients and develop resistance during infection.

**Author Contributions:** 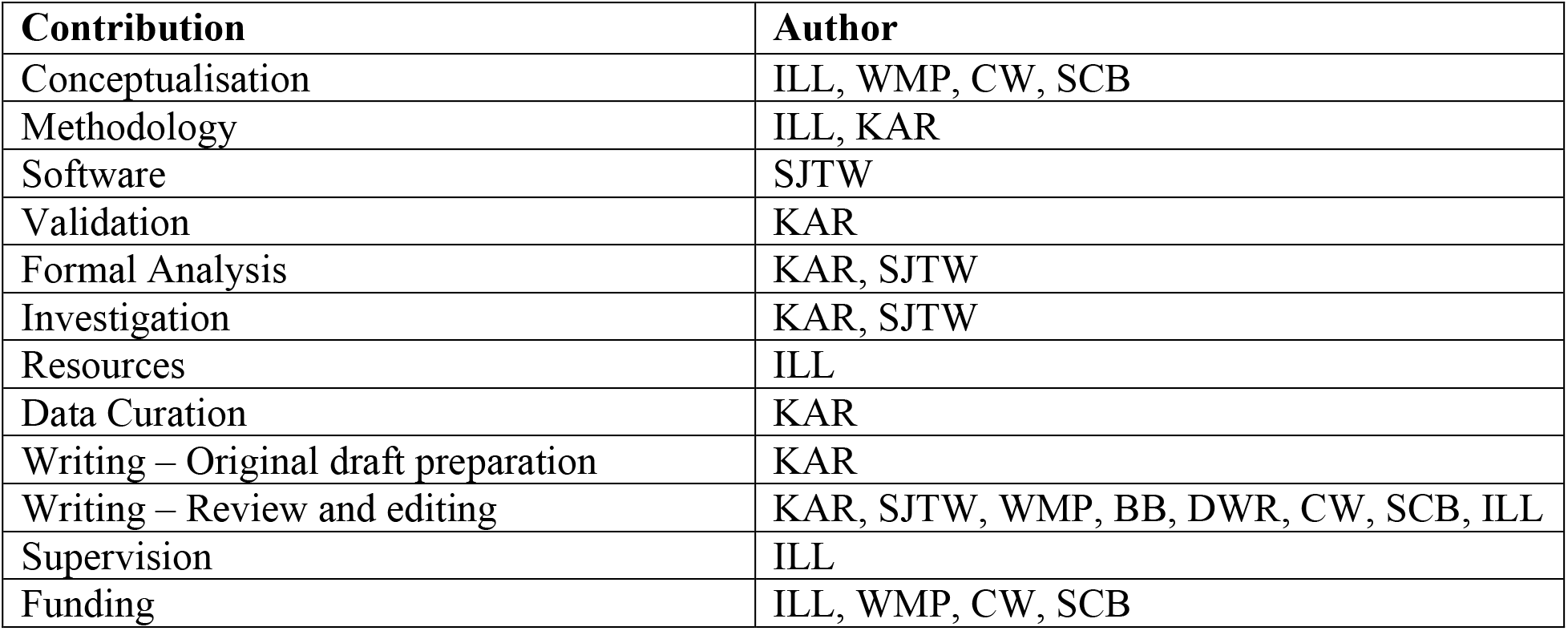

## Introduction

As an environmental bacterium, *Pseudomonas aeruginosa* has been isolated from many different niches including water sources and domestic and health-care settings (1–4). While rarely causing infections in healthy individuals, this opportunistic pathogen can cause a range of infections in people who are immunocompromised or have predisposing conditions such as cystic fibrosis (CF) (1). For adults with CF, *P. aeruginosa* is the most prevalent bacterium causing respiratory infection and *P. aeruginosa* from these infections are frequently resistant to antibiotics, complicating treatment (5). Resistance is associated with variants in key genes that reduce the intracellular concentrations of antibiotics or the affinities of target proteins for antibiotics, relative to antibiotic-susceptible isolates (6). Infection arises from exposure to environmental sources of *P. aeruginosa* included both the natural and health-care environment (1). Epidemiological studies of isolates obtained from high infection risk areas, such as health-care settings, domestic and community areas show that antibiotic resistant *P. aeruginosa* can be present providing a potential reservoir of infectious resistant bacteria (7–18). Acquisition of *P. aeruginosa* from the natural environment, typically during childhood, is also a major source of infection with a number of studies identifying genotypically indistinguishable strains in the natural environment and the respiratory tract of CF patients (3, 19–22). However antibiotic susceptibility of isolates from natural (non-man-made) environmental sources has had very limited studies and so it is not clear whether the natural environment provides a reservoir of antibiotic-resistant *P. aeruginosa* (7, 23, 24). Using a cohort of clinical isolates as a comparator, here we determined the prevalence of antibiotic resistance in *P. aeruginosa* from the natural environment (river), the domestic and community settings (swimming pool and water tank) by assessing antimicrobial susceptibility profiles and allelic variations in resistance genes of *P. aeruginosa*.

## Materials and Methods

### Isolate Selection and identification

Fifty environmental *P. aeruginosa* isolates obtained from water sources in Queensland, Australia and 42 clinical *P. aeruginosa* isolates obtained from CF patients residing in Australia and New Zealand were examined (Supplementary Table 1). Environmental isolates included those obtained from rivers (n=36), swimming pools (n=13) and one sample from a domestic water tank. Individual multi-locus sequence types (MLST) were present in singleton isolates, expect for sequence types (ST) −155, −179, −266 and −381 which were represented once each in both the CF and environmental isolate cohorts. Isolate identification and MLST typing were confirmed using molecular techniques (https://pubmlst.org/paeruginosa/) (25–27).

**Table 1:**
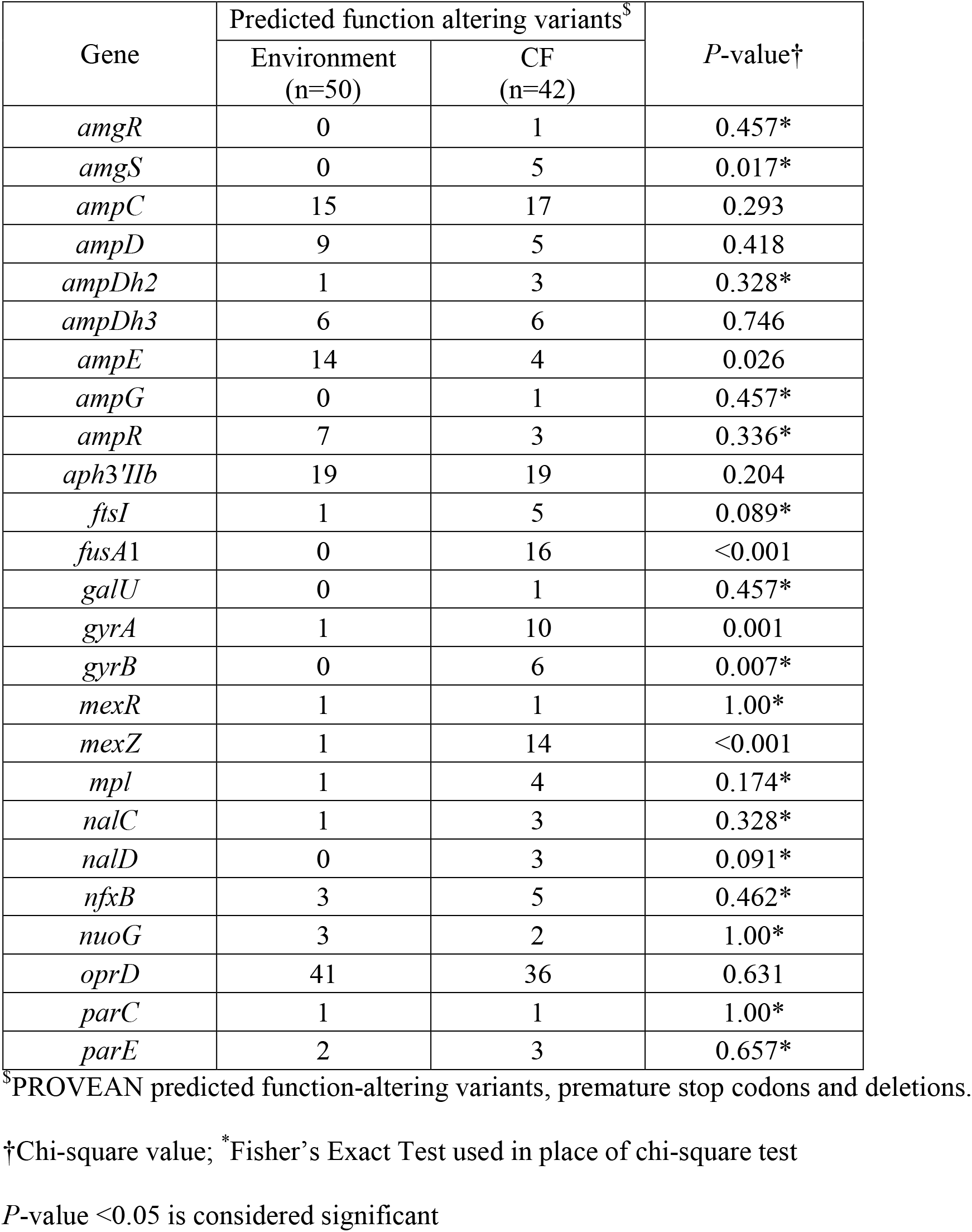
Comparison of predicted function altering variants identified in the CF and environmental isolates (n = 92)

### Antimicrobial Susceptibility Testing

The minimum inhibitory concentration (MIC) of ciprofloxacin, meropenem, tobramycin and ceftazidime was determined for all the 92 isolates using ETEST® strips and methodology. The breakpoint was determined according to the Clinical and Laboratory Standards Institute (CLSI) guidelines (https://clsi.org/).

### Resistance Gene Analysis

Whole genome sequence assemblies have been described elsewhere (28) and were generously provided by The International Pseudomonas Consortium Database (https://ipcd.ibis.ulaval.ca/). Genome assemblies were used for the comparison of 25 well characterised genes associated with antibiotic resistance, with reference sequences sourced from https://www.pseudomonas.com (Supplementary Table 2). Protein Variation Effect Analyzer (PROVEAN) version 1.1 was utilised to identify and analyse genetic variants within the chosen genes (29). Genetic variants (amino acid substitutions) were categorised in accordance to PROVEAN scores, as predicted function-altering variants (PROVEAN score: ≤ −2.5) or variants not predicted to affect function (PROVEAN score: > −2.5). Additionally, each genome was manually screened using tblastn for premature stop codons and deletions which were also classified as function-altering variants (30).

ResFinder version 3.1 was used with a prebuilt ResFinder database, including all relevant antibiotics, to determine whether resistance could be affected by acquired genes (https://cge.cbs.dtu.dk/services/ResFinder/).

### Statistical analysis

SPSS version 25 was used for statistical analysis. Pearson’s chi-square test was used to examine the association between niche and phenotype (susceptible or resistant) and niche and genotype. When more than 20% of the expected values were less than five, Fisher’s exact test were used. A *P*-value of <0.05 was considered significant.

## Results and Discussion

A total of 50 isolates of *P. aeruginosa* from the natural environment and 42 isolates from patients with CF were included in this study. The isolates are genetically diverse and broadly representative of the *P. aeruginosa* species (Supplementary Figure 1).

### Environmental isolates are antibiotic susceptible

All 50 environmental isolates were susceptible to ciprofloxacin, meropenem and tobramycin and only one of these isolates was resistant to ceftazidime (Figure 1, Supplementary Table 3). Conversely, a non-susceptible (intermediate or resistant) phenotype was observed for 15 of the 42 CF isolates. Specifically, 10%, 17% 19% and 21% of the CF isolates were non-susceptible to tobramycin, meropenem, ceftazidime and ciprofloxacin, respectively (Figure 1, Supplementary Table 4). Overall, the frequency of resistant isolates was significantly less in the environmental cohort than the CF cohort (ceftazidime *P*=0.010, ciprofloxacin *P*=0.001, meropenem *P*=0.003, tobramycin *P*=0.040). Due to limited availability of clinical information we were unable to characterise CF infections as either early/transient or chronic to assess whether resistance is more prevalent in chronically infected patients. However, when we categorised the CF isolates according to patient age (adult ≥18 years; adult n=29, paediatric n=13), isolates from two paediatric patients (15.4%) demonstrated antibiotic resistance compared with isolates from 13 adults (44.8%) (*P*=0.066) (Figure 1). Overall our findings are in agreement with a model of bacterial adaptation during infection leading to increased resistance over time (31). Our results demonstrate that environmental isolates are susceptible to clinically relevant antibiotics and that the environments we tested are not a reservoir for antibiotic resistant isolates.

**Figure 1:**
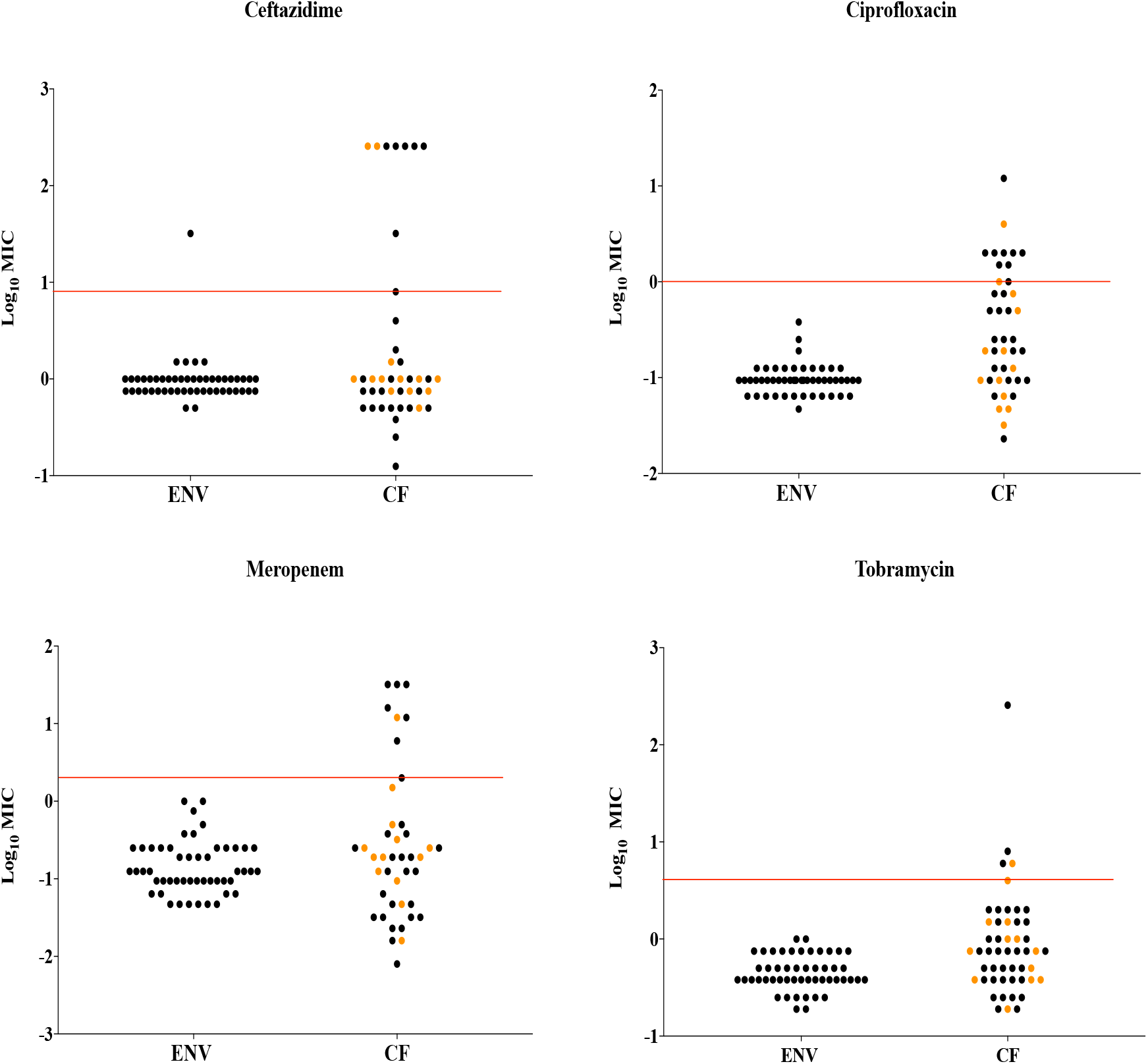
Distribution of antimicrobial susceptibility results of environmental (ENV; n=50) and cystic fibrosis (CF; n=42) isolates for ciprofloxacin, ceftazidime, meropenem and tobramycin. The horizontal red lines represent the antibiotic breakpoint as defined by the CLSI. The CF isolates have been further differentiated into samples collected from adult (black circles) and paediatric (orange circles) patients. The MIC results have been log transformed (log_10_) for ease of graphical representation.

### Environmental isolates have fewer predicted function-altering variants in resistance-associated genes

We analysed 25 well characterised resistance-associated genes for the presence of likely function-altering variants that could influence antibiotic susceptibility (Supplementary Table 2). The environmental isolates had strikingly fewer predicted function-altering variants than the CF isolates. A total of 75 predicted function-altering variants were identified within 18 of 25 resistance genes analysed for the 50 environmental isolates. No predicted function-altering variants were identified in the remaining seven genes (Supplementary Table 3). In this cohort, the predicted function-altering variants are generally not sufficient to confer antibiotic resistance as all except one isolate was fully susceptible to the antibiotics tested. In contrast, 110 predicted function-altering variants affecting all 25 resistance-associated genes were identified in the 42 CF isolates. Function-altering variants not analysed by PROVEAN, including premature stop-codons and deletions, were absent in the environmental cohort but were present in nine CF isolates (premature stop-codons n=8, deletions n=2) (Supplementary Table 4). Overall, variants in the environmental cohort had a significantly higher PROVEAN score than those in the CF cohort (*P*<0.001), indicating the presence of fewer amino acid substitutions that may affect function of resistance-associated proteins (Supplementary Figure 2).

The frequency of function-altering variants within each gene was determined for both cohorts of isolates (Table 1). Statistical differences were noted for 6 genes in the CF cohort with a greater number of function-altering variants being present in *amgS* and *fusA*1 (tobramycin resistance), *gyrA* and *gyrB* (ciprofloxacin resistance) and *mexZ* (broad-spectrum resistance) genes for isolates in the CF cohort, and in *ampE* (β-lactam resistance) for the natural environmental isolates (Table 1). Specific variants present in these genes, such as T83I (*gyrA*), and R504C (*ftsI*) previously associated with resistance were only present in the CF isolate cohort only (32, 33). However, a strong association between individual gene variants and resistance phenotype was not observed (Supplementary Table 4) due to the low numbers of isolates with variants for each gene combined with the multifactorial nature of antibiotic resistance in *P. aeruginosa*.

ResFinder analysis assessing the presence of horizontally acquired resistance genes identified a gene *crpP* associated with fluoroquinolone resistance (34). However there was no correlation between ciprofloxacin resistance and the presence of *crpP* for the isolates in this study (Supplementary Table 3 and 4). No other horizontally-transferred resistance genes were identified.

In conclusion, our findings show that isolates from *P. aeruginosa* from natural environments have low frequencies of antibiotic resistance, and of genetic variants associated with resistance, compared to isolates from patients with CF. These findings indicate that the natural environment is unlikely to act as a reservoir of antibiotic-resistant *P. aeruginosa*. They are consistent with a model in which patients are infected by antibiotic-sensitive *P. aeruginosa* from the environment which then evolves to become antibiotic-resistant during infection (31).

## Acknowledgements

We would like to acknowledge and thank the individuals with cystic fibrosis for providing samples. We thank the members of the CF healthcare teams at The Prince Charles Hospital, Lady Cilento Children ‘s Hospital, Royal Hobart Hospital, Royal Children’s Hospital, Melbourne and Westmead Children ‘s Hospital, Australia and Dunedin Public Hospital, New Zealand and the staff of the various microbiology departments for their ongoing support of research. We thank Dr Timothy Kidd, Professor Claire Wainwright and Professor Keith Grimwood for access to the Australian isolate collection. We thank Professor Roger C. Lévesque and colleagues at the Institute of Integrative Biology and Systems at Université Laval, Québec for providing whole genome sequence assemblies of isolates analysed in this study.

## Transparency declarations

K.A.R, S.J.T-W, W.M.P, B.B, D.W.R and I.L.L have nothing to declare. S.C.B is a Member of Advisory Board, Member of Writing group, Site Principle Investigator, author on several Rempex sponsored studies, a member of advisory boards for Vertex, Abbvie and Galapagos and has received support to attend meetings including advisory boards and investigator meetings. CW was a member of a Chiesi Limited CF Microbiology Advisory Board (12 September 2018).

