## Supplementary Tables 1_2 and Supplementary Figures 1_2 for "Genomic and phenotypic comparison of environmental and patient-derived isolates of *Pseudomonas aeruginosa* suggest that antimicrobial resistance is rare within the environment"

Supplementary Table 1: List of the source, sampling details and multi-locus sequence typing (MLST) data for all *Pseudomonas aeruginosa* isolates included in this study (n=92).

| Sequence Identifier | Source | MLST type | Patient Location/Sample Information | Isolate ID from IPCD* | Accession Number | Isolate Reference |
|---|---|---|---|---|---|---|
| AUS110 | Environment | 826 | Brisbane River | 357 | SAMN10478427 | 2, 8 |
| AUS116 | Environment | 832 | Brisbane River | 363 | In progress | 2 |
| AUS122 | Environment | 838 | Brisbane River | 367 | SAMN10478429 | 2, 8 |
| AUS125 | Environment | 841 | Municipal Pool | 369 | SAMN10478430 | 2, 8 |
| AUS128 | Environment | 843 | Brisbane River | 371 | SAMN10478431 | 2, 8 |
| AUS136 | Environment | 850 | Brisbane River | 377 | SAMN10478432 | 2, 8 |
| AUS139 | Environment | 853 | Municipal Pool | 380 | SAMN10478433 | 2, 8 |
| AUS141 | Environment | 855 | Brisbane River | 382 | SAMN10478435 | 2, 8 |
| AUS153 | Environment | 867 | Brisbane River | 392 | SAMN10478436 | 2, 8 |
| AUS155 | Environment | 869 | Brisbane River | 394 | SAMN10478437 | 2, 8 |
| AUS156 | Environment | 870 | Brisbane River | 395 | SAMN10478438 | 2, 8 |
| AUS157 | Environment | 871 | Brisbane River | 396 | SAMN10478439 | 2, 8 |
| AUS158 | Environment | 872 | Municipal Pool | 397 | SAMN10478440 | 2, 8 |
| AUS174 | Environment | 888 | Brisbane River | 411 | SAMN10478441 | 2, 8 |
| AUS175 | Environment | 889 | Brisbane River | 412 | SAMN10478442 | 2, 8 |
| AUS176 | Environment | 890 | Municipal Pool | 413 | SAMN10478443 | 2, 8 |
| AUS177 | Environment | 891 | Municipal Pool | 414 | SAMN10478444 | 2, 8 |
| AUS178 | Environment | 892 | Municipal Pool | 415 | SAMN10478445 | 2, 8 |
| AUS195 | Environment | 115 | Municipal Pool | 215 | SAMN07423914 | 2, 7 |
| AUS209 | Environment | 161 | Brisbane River | 229 | SAMN10478413 | 2, 8 |
| AUS214 | Environment | 172 | Domestic Pool | 236 | SAMN10478414 | 2, 8 |
| AUS217 | Environment | 191 | Municipal Pool | 241 | SAMN07423922 | 2, 7, 8 |
| AUS221 | Environment | 205 | Brisbane River | 245 | SAMN10478415 | 2, 8 |
| AUS222 | Environment | 209 | Brisbane River | 247 | SAMN10478417 | 2, 8 |
| AUS226 | Environment | 215 | Brisbane River | 252 | SAMN10478419 | 2, 8 |
| AUS227 | Environment | 216 | Brisbane River | 253 | SAMN10478420 | 2, 8 |
| AUS258 | Environment | 212 | Municipal Pool | 249 | SAMN10478418 | 2, 8 |
| AUS265 | Environment | 645 | Municipal Pool | 325 | SAMN10478425 | 2, 8 |
| AUS276 | Environment | 9 | Brisbane River | 204 | In progress | 2 |
| AUS277 | Environment | 564 | Domestic Water Tank | 317 | SAMN10478423 | 2, 8 |
| AUS306 | Environment | 381 | Brisbane River | 294 | SAMN07423940 | 2, 7 |
| AUS311 | Environment | 854 | Brisbane River | 381 | SAMN10478434 | 2, 8 |
| AUS321 | Environment | 316 | Municipal Pool | 289 | SAMN07423938 | 2, 7 |
| AUS328 | Environment | 385 | Brisbane River | 295 | SAMN07423941 | 2, 7 |
| AUS339 | Environment | 360 | Brisbane River | 292 | SAMN07423939 | 2, 7 |
| AUS343 | Environment | 266 | Brisbane River | 274 | SAMN07423931 | 2, 7 |
| AUS355 | Environment | 553 | Brisbane River | 312 | SAMN07423947 | 2, 7 |
| AUS437 | Environment | 155 | Brisbane River | 473 | In progress | 2 |
| AUS449 | Environment | 930 | Brisbane River | 451 | SAMN10478454 | 2, 8 |
| AUS452 | Environment | 257 | Brisbane River | 270 | SAMN10478421 | 2, 8 |
| AUS464 | Environment | 909 | Municipal Pool | 432 | SAMN10478446 | 2, 8 |
| AUS465 | Environment | 147 | Brisbane River | 227 | SAMN07423917 | 2, 7 |
| AUS502 | Environment | 179 | Logan River | 239 | SAMN07423921 | 2, 7 |
| AUS503 | Environment | 810 | Logan River | 353 | SAMN07423960 | 2, 7 |
| AUS504 | Environment | 389 | Logan River | 297 | SAMN07423942 | 2, 7 |
| AUS505 | Environment | 923 | Logan River | 445 | SAMN10478449 | 2, 8 |
| AUS506 | Environment | 863 | Logan River | 389 | SAMN07423970 | 2, 7 |
| AUS507 | Environment | 924 | Brisbane River | 446 | SAMN07423983 | 2, 7 |
| AUS510 | Environment | 925 | Brisbane River | 447 | SAMN10478450 | 2, 8 |
| AUS511 | Environment | 926 | Brisbane River | 448 | SAMN10478451 | 2, 8 |

| | | | | | | |
|---|---|---|---|---|---|---|
| AUS021 | Adult CF | 266 | Brisbane | 494 | SAMN07424000 | 1, 7 |
| AUS026 | Paediatric CF | 777 | Brisbane | 334 | SAMN07423951 | 1, 7 |
| AUS029 | Adult CF | 779 | Brisbane | 336 | SAMN07423952 | 1, 7 |
| AUS050 | Paediatric CF | 155 | Brisbane | 470 | SAMN07423990 | 1, 7 |
| AUS054 | Adult CF | 788 | Brisbane | 340 | SAMN07423954 | 1, 7 |
| AUS058 | Adult CF | 494 | Brisbane | 306 | SAMN07423944 | 1, 7 |
| AUS066 | Adult CF | 798 | Brisbane | 345 | SAMN07423955 | 1, 7 |
| AUS073 | Adult CF | 800 | Brisbane | 346 | SAMN07423956 | 1, 7 |
| AUS077 | Paediatric CF | 801 | Brisbane | 347 | SAMN07423957 | 1, 7 |
| AUS083 | Adult CF | 259 | Brisbane | 271 | SAMN07423929 | 1, 7 |
| AUS088 | Adult CF | 262 | Brisbane | 273 | SAMN07423930 | 1, 7 |
| AUS089 | Adult CF | 805 | Brisbane | 349 | SAMN07423958 | 1, 7 |
| AUS093 | Adult CF | 809 | Brisbane | 352 | SAMN07423959 | 1, 7 |
| AUS105 | Paediatric CF | 821 | Brisbane | 354 | SAMN07423961 | 1, 7 |
| AUS106 | Adult CF | 822 | Brisbane | 355 | SAMN07423962 | 1, 7 |
| AUS113 | Paediatric CF | 829 | Brisbane | 360 | In progress | 1 |
| AUS127 | Paediatric CF | 2 | Brisbane | 197 | SAMN07423909 | 1, 7 |
| AUS148 | Adult CF | 862 | Brisbane | 388 | SAMN07423969 | 1, 7 |
| AUS159 | Adult CF | 873 | Brisbane | 398 | In progress | 1 |
| AUS304 | Adult CF | 655 | Brisbane | 327 | SAMN07423950 | 1, 7 |
| AUS305 | Adult CF | 381 | Brisbane | 506 | SAMN07424003 | 1, 7 |
| AUS327 | Adult CF | 443 | Brisbane | 300 | SAMN07423943 | 1, 7 |
| AUS476 | Adult CF | 273 | Brisbane | 277 | SAMN07423933 | 1, 7 |
| AUS537 | Paediatric CF | 27 | Melbourne | 468 | SAMN07423989 | 1, 7 |
| AUS538 | Paediatric CF | 179 | Melbourne | 485 | In progress | 1 |
| AUS631 | Paediatric CF | 1099 | Brisbane | 460 | SAMN07423986 | 3, 7 |
| AUS674 | Paediatric CF | 1102 | Sydney | 462 | SAMN07423987 | 3, 7 |
| AUS675 | Paediatric CF | 1103 | Sydney | 463 | SAMN07423988 | 3, 7 |
| AUS702 | Paediatric CF | 782 | Melbourne | 399 | SAMN07423953 | 1, 7 |
| AUS717 | Adult CF | 775 | Brisbane | 532 | SAMN10478459 | 1 |
| DUN-001C | Adult CF | 244 | Dunedin | 980 | SAMN07424118 | This study, 7 |
| DUN-002C | Adult CF | 274 | Dunedin | 991 | SAMN07424127 | This study, 7 |
| DUN-003B | Adult CF | 189 | Dunedin | 995 | SAMN07424128 | This study, 7 |
| DUN-004 | Adult CF | 224 | Dunedin | 1001 | SAMN07424129 | This study, 7 |
| DUN-009B | Adult CF | 395 | Dunedin | 1014 | SAMN07424133 | This study, 7 |
| DUN-014-2A | Adult CF | 1225 | Dunedin | 1020 | SAMN07424136 | This study |
| DUN-015A | Adult CF | 499 | Dunedin | 1027 | SAMN07424137 | This study, 7 |
| S2820 | Adult CF | 110 | Dunedin | 905 | SAMN07424099 | This study, 7 |
| U0306 | Paediatric CF | 781 | Hobart | 934 | SAMN07424106 | This study, 7 |
| U0310A | Adult CF | 649 | Hobart | 935 | SAMN07424107 | This study, 7 |
| U413A | Adult CF | 988 | Hobart | 962 | SAMN07424112 | This study, 7 |
| U451 | Adult CF | 242 | Hobart | 976 | SAMN07424115 | This study, 7 |
| PAO1 | NA | 549 | NA | NA | NC_002516.2 | 4, 5 |
| UCBPP-PA14 | NA | 253 | NA | NA | NC_008463 | 4, 6 |

*International Pseudomonas Consortium Database https://ipcd.ibis.ulaval.ca/index

Supplementary Table 2: Key antimicrobial resistance genes examined for PROVEAN predicted function altering variants.

| Gene | Locus tag | Reference |
|---|---|---|
| *amgR* | PA5200 | 1-4 |
| *amgS* | PA5199 | 1-5 |
| *aph3'IIb* | PA4119 | 6 |
| *ampC* | PA4110 | 5, 7, 8 |
| *ampD* | PA4522 | 4, 5, 8 |
| *ampDh*2 | PA5485 | 5, 8 |
| *ampDh*3 | PA0807 | 5, 8 |
| *ampE* | PA4521 | 8, 9 |
| *ampG* | PA4393 | 7, 8 |
| *ampR* | PA4109 | 5, 8 |
| *ftsI* | PA4418 | 5, 9 |
| *fusA*1 | PA4266 | 4, 5 |
| *galU* | PA2023 | 7, 10 |
| *gyrA* | PA3168 | 4, 5, 7, 11 |
| *gyrB* | PA0004 | 4, 5, 7, 11 |
| *mexR* | PA0424 | 4, 5, 7, 11 |
| *mexZ* | PA2020 | 4, 5, 7, 11 |
| *mpl* | PA4020 | 4, 9 |
| *nalC* | PA3721 | 5 |
| *nalD* | PA3574 | 5 |
| *nfxB* | PA4600 | 5, 7 |
| *nuoG* | PA2642 | 7, 10 |
| *oprD* | PA0958 | 4, 5 |
| *parC* | PA4964 | 5, 7, 9, 11 |
| *parE* | PA4967 | 5, 7, 11 |

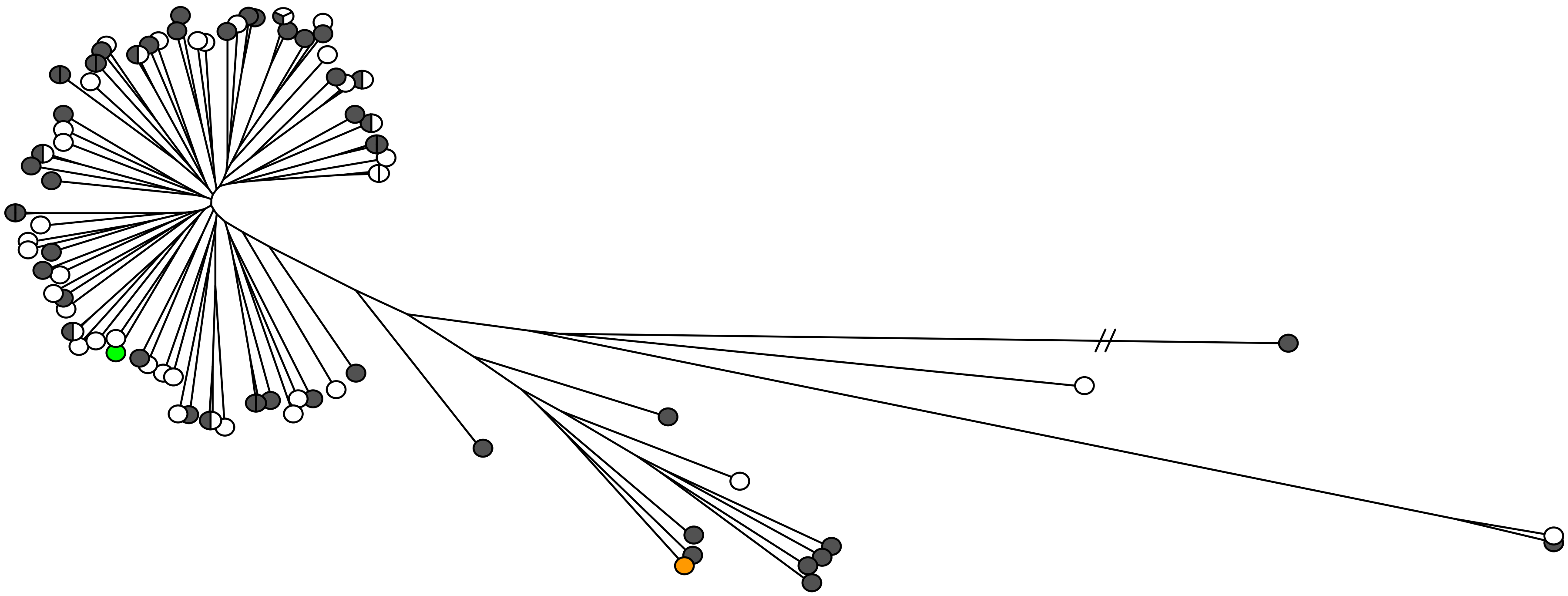

Supplementary Figure 1: Phylogenetic tree displaying the genotypic diversity of the environmental and CF isolates (n = 92) based on whole genome sequence data. The tree was constructed with parSNP version 1.2, using *P. aeruginosa* PAO1 and PA14 as reference strains. This generated a core genome comparison of 56% (~3.5 mbp) and was visualised as a tree using FigTree version 1.4.3 (http://tree.bio.ed.ac.uk/software/figtree). Grey circles represent isolates from the natural environment; white circles represent CF isolates; green and orange circles represent the reference strains PAO1 and UCBPP-PA14, respectively.

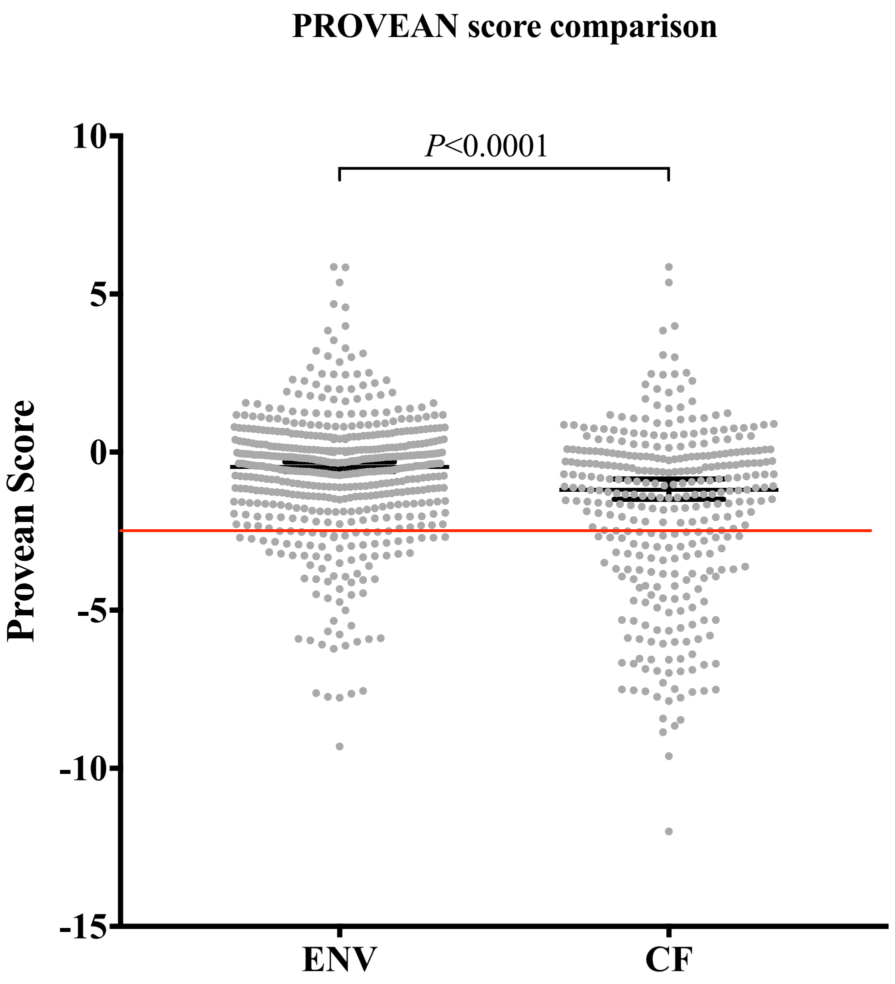


Supplementary Figure 2: Comparison of the PROVEAN scores of antibiotic resistance-associated genes in environmental and CF isolates of *P. aeruginosa*. Each grey dot represents a protein variant. The red line represents the cut-off between predicted function altering variants (≤ -2.5) and variants not predicted to affect function (> -2.5). The error bars represent the median and 95% confidence interval.
